# Inkjet-printed graphene multielectrode arrays: an accessible platform for *in vitro* cardiac electrophysiology

**DOI:** 10.1101/2024.09.09.611887

**Authors:** Jairo Lumpuy-Castillo, Yujie Fu, Alan Ávila, Kateryna Solodka, Jiantong Li, Oscar Lorenzo, Erica Zeglio, Leonardo D. Garma

**Affiliations:** Laboratory of Diabetes and Vascular Pathology, IIS-Fundación Jiménez Díaz (CIBERDEM), Universidad Autónoma, 28040 Madrid, Spain; School of Electrical Engineering and Computer Science, KTH Royal Institute of Technology, Electrum 229, Kista, 16440 Sweden; Division of Nanobiotechnology, Department of Protein Science, Science for Life Laboratory, School of Engineering Sciences in Chemistry, Biotechnology and Health, KTH Royal Institute of Technology, Solna, 171 65 Sweden; AIMES-Center for the Advancement of Integrated Medical and Engineering Sciences, Department of Neuroscience, Karolinska Institute, Solna, 171 65 Sweden; Wallenberg Initiative Materials Science for Sustainability, Department of Materials and Environmental Chemistry, Stockholm University, Stockholm, 114 18 Sweden; Digital Futures, Stockholm, SE-100 44 Sweden; Breast Cancer Clinical Research Unit, Centro Nacional de Investigaciones Oncológicas – CNIO, Madrid, Spain

## Abstract

*In vitro* models have now become a realistic alternative to animal models for cardiotoxicity assessment. However, the cost and expertise required to implement *in vitro* electrophysiology systems to study cardiac cells poses a strong obstacle to their widespread use. This study presents a novel, cost-effective approach for *in vitro* cardiac electrophysiology using fully-printed graphene-based microelectrode arrays (pGMEAs) coupled with an open-source signal acquisition system. We characterized the pGMEAs’ electrical properties and biocompatibility, observing low impedance values and cell viability. We demonstrated the platform’s capability to record spontaneous electrophysiological activity from HL-1 cell cultures, and we monitored and quantified their responses to chemical stimulation with noradrenaline. This study demonstrates the feasibility of producing fully-printed, graphene-based devices for *in vitro* electrophysiology. The accessible and versatile platform we present here represents a step further in the development of alternative methods for cardiac safety screening.

## Introduction

Assessing cardiac safety is a crucial step during drug development, as cardio-related adverse reactions can have a dramatic impact on patient’s life, leading to severe health problems or even death^1,2^. Drug-derived cardiotoxicity is generally evaluated using animal models. However, because of wide interspecies variability, especially at the cardiovascular level, animal models frequently fail to predict the therapeutic response in humans^3–5^. Moreover, the use of animal models is laborious and expensive, as well as ethically questionable. Aiming for the reduction and replacement of animal models, *in vitro* systems are emerging as a potential alternative for cardiac safety evaluation^6,7^. *In vitro* models allow for the direct recording of electrophysiological signals from electrically active cells, such as cardiomyocytes, providing a more accurate perspective of human physiology and making them an attractive tool for cardiac monitoring.

Because of its high temporal resolution and the high signal-to-noise ratio (SNR) obtained, the patch-clamp technique is referred to as the gold standard for intracellular electrophysiological recordings^8,9^. However, several limitations are associated with this technique. First, the lack of spatial resolution, which restricts the recordings to single-cell level and, second, the high invasiveness, resulting from the disruption of the cell membrane and decreased electrical activity. As a consequence, it is not possible to perform long-term recordings, lasting several hours or days^10,11^. Optical imaging techniques, employing fluorescent voltage- or calcium-sensitive dyes, are also a powerful tool for electrophysiology studies. Optical techniques allow to record cardiac signals with high spatial, allowing to study the whole cell population^12,13^. However, optical imaging suffers from several drawbacks, such as low SNR, photobleaching, phototoxicity, and dye internalization, which strongly limit their use^14^. To overcome these challenges, microelectrode arrays (MEAs) emerged as an attractive platform for *in vitro* electrophysiology studies^15,16^. MEAs allow for the continuous, non-invasive, long-term recording of electrophysiological signals, providing high spatial and temporal resolution^17–21^. Nevertheless, despite their many advantages, the use of MEA-based platforms is still limited. This is primarily due to the relatively low SNR obtained with respect to path-clamp, and the high costs related to the materials and fabrication process (typically gold and photolithography, respectively). Furthermore, MEAs require costly signal acquisition hardware, which usually relies on closed-source software. Although some advances have been made in the development of cost-effective open-source solutions, there is currently a very limited number of designs available, which are either bulky and reliant on obsolete components^22^, or not adapted for planar MEAs and with limited throughput^23^.

The advances in printed electronics have now created a realistic alternative to the costlier traditional microfabrication strategies for the production of bioelectronic devices. Printing technologies are highly versatile and enable the creation of novel architectures with high resolution and reproducibility, favoring the production of large-scale, customizable, and cost-effective devices for biomedical applications^24–29^. Furthermore, the recent progress in materials chemistry has led to the development of novel materials with outstanding bioelectronic properties, such as conductive polymers and carbon-based materials. These materials constitute promising alternatives to the metallic electrodes employed in conventional MEA devices^30–34^. Among these, graphene, a 2D carbon-based material, exhibits a unique combination of electrical, optical, and mechanical properties. Graphene possesses high electron mobility, in the order of 10^5^ cm^2^/Vs^35,36^, making it one of the most conductive materials (at room temperature). In addition, graphene is optically transparent^37,38^, and presents exceptional mechanical strength^39^. Furthermore, its flexibility^40–42^ and biocompatibility^34,43–45^ make graphene a very attractive material to serve as the interface between electronic systems and the biological world. Due to these extraordinary properties, graphene and its derivatives have been widely explored for a broad range of biomedical applications, including biosensing^46,47^, drug delivery^48,49^, and tissue engineering^50,51^. Graphene-based MEAs (GMEAs) capable of recording the electrophysiological activity from cardiac cells have been developed in the past years^34,52–54^, highlighting the potential of these devices as a valuable platform for electrophysiological signal investigation. Typically, GMEAs fabrication relies on multi-step microfabrication techniques.^55–59^ Although recently the fabrication of printed GMEAs (pGMEAs) has been reported, so far pGMEAs have only been used for cell survival monitoring via electrochemical impedance spectroscopy, with no examples of devices tested for *in vitro* cardiac electrophysiology^60,61^.

Here, we introduce a fully-printed microelectrode array based on graphene electrodes, together with a simple, open-source signal acquisition system, conforming a highly accessible *in vitro* platform for electrophysiological studies. We characterized the electrical properties and the biocompatibility of pGMEAs, observing impedance values as low as 19.2 kΩ at 1 kHz and high cell viability. We then used our acquisition system to record the spontaneous electrophysiological activity of HL-1 cell cultures on pGMEAs and commercial MEAs. Finally, we demonstrated the capability of our approach to monitor and quantify the response of cardiac cell cultures to chemical stimulation using noradrenaline.

## Results

### Fabrication of pGMEAs

We designed an MEA with 16 sensing electrodes and a reference electrode. The fabrication strategy included the deposition of three different layers via inkjet printing on a glass substrate. A first conductive layer, defining the external connection pads and with tracks leading to the center of the MEA, was printed using silver ink. A second layer was printed at the center of the device using a graphene ink, which was rendered conductive after annealing at 350 ºC (see experimental section for details). The same graphene ink (a 10:1 ethyl cellulose:graphene solution) was used to print a passivation layer without annealing, as the material is not conductive unless the high temperature process is applied. The passivation material covered both conductive layers, exposing only the external connectors, the ends of the tracks in the center of the device through 16 square openings of 400×400 μm for the microelectrodes and a triangular section of 12.5 mm^2^ of the conductive graphene layer to be used as the reference electrode (Figure 1A,B, Supplementary Data 1).

**Figure 1.**
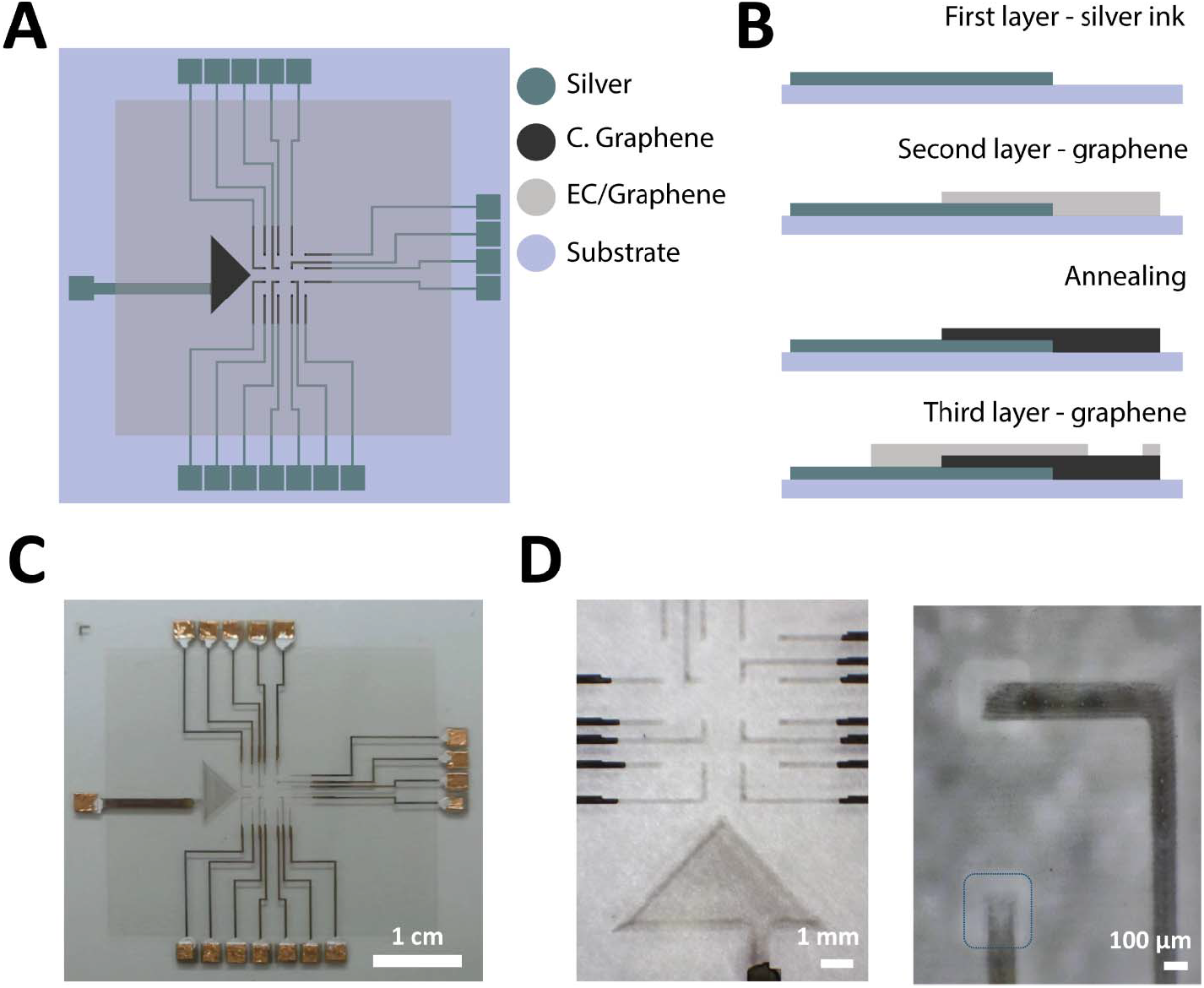
**A**. Layout of the pGMEA, with elements colored by the corresponding material: silver ink, conductive graphene, ethyl cellulose(EC)/graphene (non-conductive), and the glass substrate. **B**. Schematic of the printed electrodes fabrication strategy, with materials colored as in panel A. **C**. A finished pGMEA, with the contact pads covered in copper tape. **D**. Microscopy image of the electrodes on a pGMEA (left) and a close-up of two electrodes (right). The blue line highlights an opening on the passivation layer.

After the printing process was complete, the external connectors of the pGMEAs (contact pads) were covered with silver paste and copper tape to avoid damaging the printed material underneath when the devices were in use. The fabricated pGMEA devices reliably reproduced the initial design (Figure 1C), and its main features could be directly observed under the microscope (Figure 1D).

### Characterization of pGMEAs

We measured the size of the printed graphene electrodes and the openings on the passivation layer around them. The effective dimensions of the electrodes presented only slight deviations from the original design: the openings on the passivation layer were on average 387 µm (σ = 21 µm, N = 5) x 393 µm (σ = 27 µm, N = 5) instead of the intended 400×400 µm, and the conductive tracks exposed had an average width of 235 µm (σ = 15 µm, N = 5) rather than the expected 240 µm. The dimensions observed on the pGMEAs were remarkably consistent, with a standard deviation below 30 µm in all cases (Figure 2A, Supplementary Figure 1). The length of the exposed conductive tracks was on average 283 µm (σ = 22 µm, N = 5), thus making the average effective electrode area on the pGMEAs 66,505 µm^2^.

**Figure 2.**
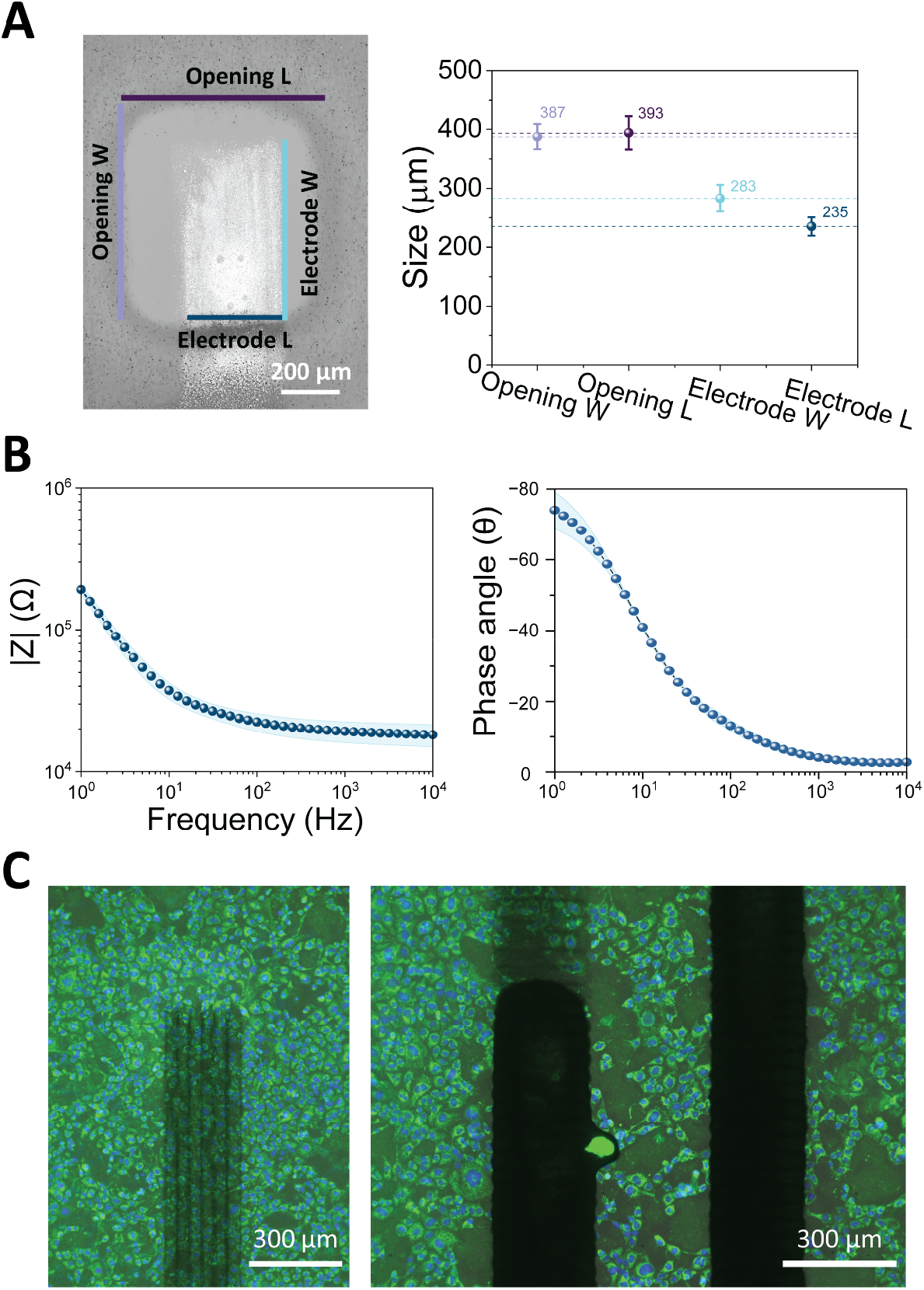
**A**. Left: Electrode and opening measures overlayed on a microscopy image of a printed graphene electrode. Right: mean values observed for each of the four measures on 5 electrodes. The whiskers indicate the standard deviation of the measures. **B**. Impedance spectrum of 5 printed graphene electrodes from a pGMEA. The left panel shows the observed magnitude of the impedance at difference frequencies. The right panel shows the phase angle diagram. In both panels, the dots represent the mean value observed across all the measured electrodes (n = 5), while the light blue trend line represents the standard deviation values from the samples both above and below the mean **C**. Staining of HL-1 cells plated on pGMEA. The image overlays the intensities of MitoTracker Green (green) and Hoechst 33342 (blue) stainings with the brightfield signal, which mark live cells and nuclei respectively.

The electrical properties of the printed graphene electrodes were characterized by electrochemical impedance spectroscopy (EIS). EIS experiments were performed in a three-electrode setup, using 1X PBS as electrolyte. Electrode impedance was examined across a frequency spectrum spanning from 1 to 10^5^ Hz, encompassing the relevant range for electrophysiological signal detection. The Bode plot shows the typical response observed for the printed graphene electrodes, with a linear impedance decrease at low frequencies, up to 10 Hz, followed by a nearly flat behavior at higher frequencies. The phase spectrum shows a predominantly capacitive behavior of the printed graphene electrodes at low frequencies, becoming nearly resistive at higher frequencies (phase near 0°). The observed impedance at 1 kHz presented values of 19.2 kΩ (σ = 3.2 kΩ, N = 5, normalized value = 2.5 Ωcm^2^), matching the impedance values previously reported for PEDOT:PSS^32,62,63^ (Figure 2B).

To verify that the pGMEAs were a suitable substrate for cell culture, we plated them with cardiomyocyte-like HL-1 cells, a cell line derived from the atrial tissue of transgenic mice^64^. After maintaining the cells in culture for 48 hours, we used a combination of MitoTracker Green™ (MTG) and Hoechst 33342, which selectively stain mitochondria in live cells and nuclei, respectively. Using this combination, live cells could be identified as those presenting a regular, round nucleus stained in blue surrounded by a cell body stained in green.

The staining revealed that the cells proliferated normally on the surface of the devices, as a nearly confluent layer of cells could be observed on electrodes, the passivated tracks, and the reference electrode (Figure 2C, Supplementary Figure 2).This seems to indicate that none of the materials used inhibited cell proliferation nor induced cell death.

### Acquisition system

To be able to monitor and record electrical signals from the electrodes on the pGMEAs, we designed and produced a custom signal acquisition system. This system consisted of two main components connected via a single HDMI cable: 1) a printed circuit board (PCB) with 2×32-channel amplifiers (INTAN Technologies, USA) onto which we mounted an MEA connector board and 2) a field programmable gate array (FPGA, Opal Kelly, USA) mounted on a breakout board (Figure 3A, B). This design intends to reduce the amount of electronics around the MEAs, so that the amplifier and MEA connector boards can be kept under a cell culture hood or inside an incubator, maintaining optimal cell culture conditions while recording. The MEA connector board was mounted on the amplifier board using two arrays of vertical 2.54 mm pins, so that it would be easily replaceable by a board adapted to different MEA designs or layouts. The amplifier board allowed to ground the reference of the amplifiers and/or to connect it to the internal reference of the MEAs. The PCB schematics and the list of components are provided in the supplementary materials (Supplementary Data 2).

**Figure 3.**
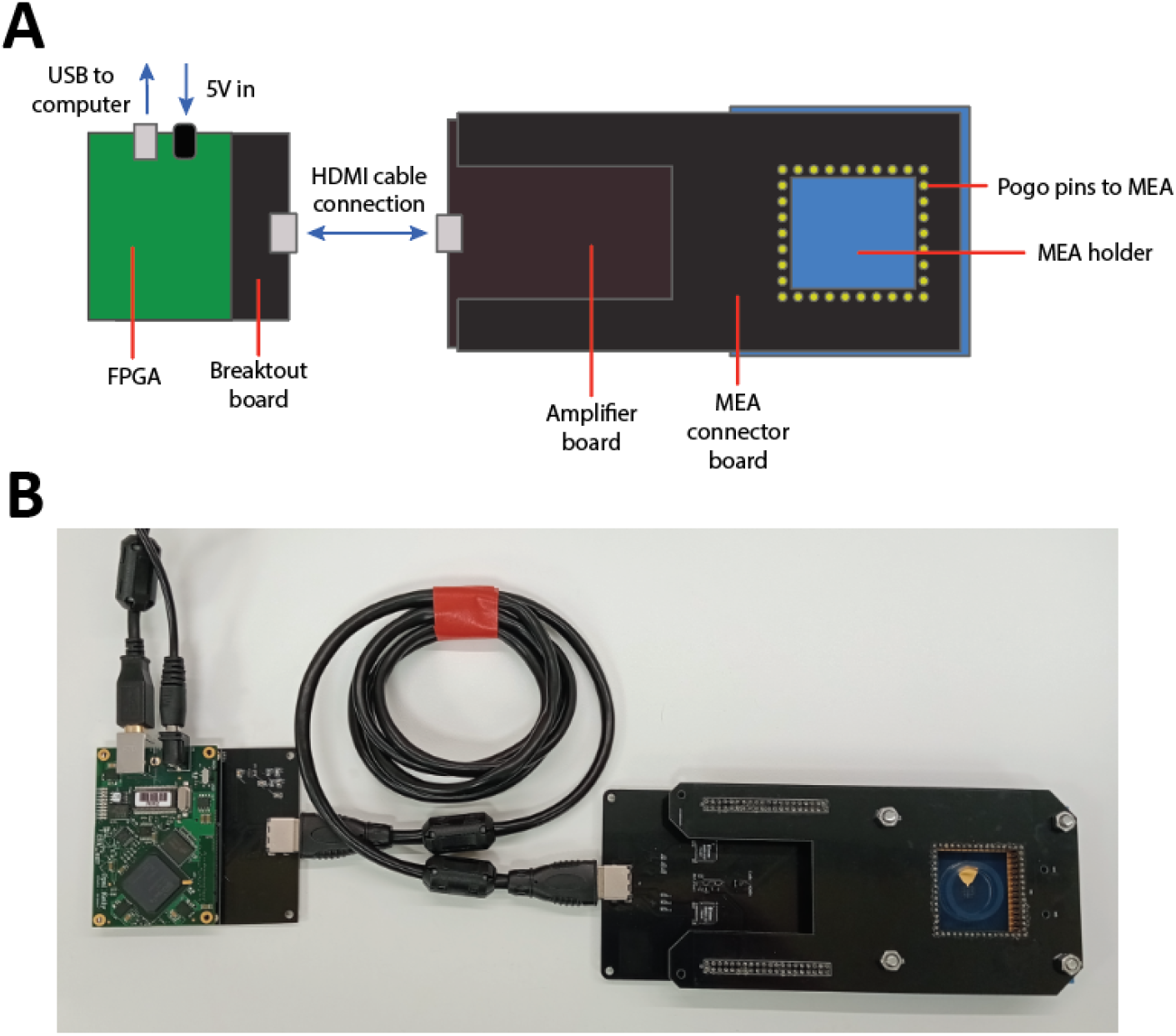
**A**. Schematic of the components and connections of the acquisition system **B**. The assembled acquisition system connected to a commercial MEA.

### Cardiac electrophysiology

The main intended application of the pGMEAs is to monitor the electrophysiological activity of cardia cell cultures. Therefore, we used them to record the spontaneous electrical activity of HL-1 cell cultures and compare it with signals obtained using commercial MEAs with flat gold electrodes.

We recorded the spontaneous electrophysiological activity from HL-1 cultures on 2 pGMEAs and on one commercial MEA, observing voltage traces with local-field potential (LFP)-like signals in all three devices (Figure 4A). We detected regular LFPs in 12 pGMEAs electrodes and on 40 gold electrodes. We analyzed 90-second-long recordings and observed that the amplitude of the signals was significantly lower on the pGMEAs, with an average of 58.65 µV (σ = 32.87 µV, N = 12) compared to the 214.86 µV (σ = 137.64 µV, N = 45) of the commercial MEAs. This difference in amplitude could be explained by the differences in electrode size, as the electrodes on the pGMEAs were almost one order of magnitude larger than the ones on the commercial devices (66,505 µm^2^ on the pGMEAs compared to 7,850 µm^2^ on the regular MEAs). The larger electrode size leads to more spatial averaging, reducing the peak amplitudes^65^. The SNR was likewise significantly higher on the commercial devices: we obtained an average value of 11.41 dB (σ = 3.82 dB, N = 12) for the pGMEAs electrodes and a value of 14.91 dB (σ = 4.37 dB, N = 45) for the gold electrodes on the commercial devices (Figure 4B).

**Figure 4.**
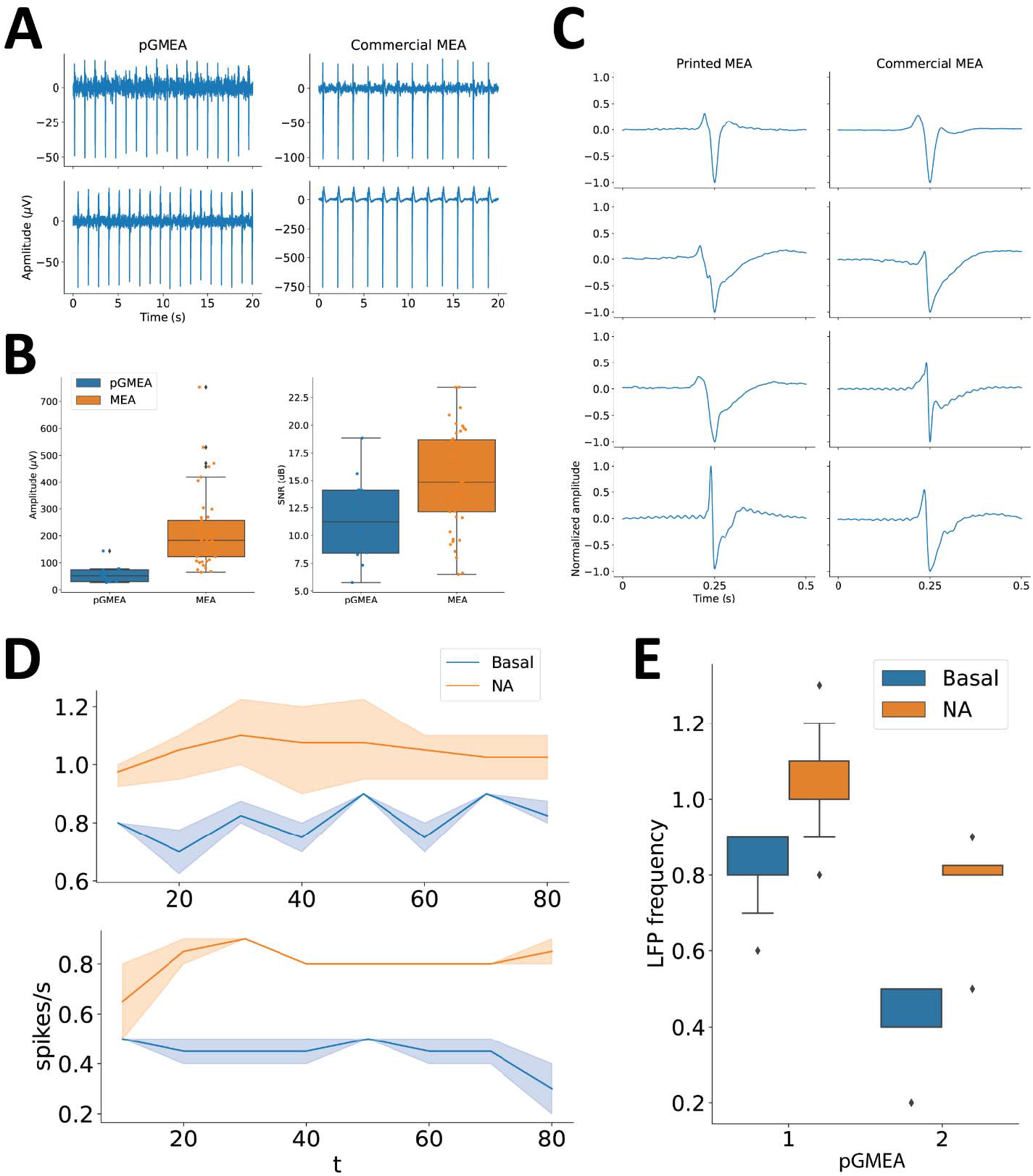
**A**. Voltage traces recorded from two different electrodes on a pGMEA (left) and on a commercial MEA (right) **B**. Distribution of peak amplitudes (left) and signal SNR (right) on 90-second recordings from pGMEAs (N = 12 electrodes) and commercial MEAs (N = 40 electrodes). **C**. Mean waveforms of the spikes detected on the recordings obtained from four different electrodes on a pGMEA (left) and a commercial MEA (right), normalized by amplitude. **D**. Firing rates detected on two separate pGMEAs before (blue) and after (orange) stimulation with noradrenaline (NA) (N = 2 electrodes per device), calculated every 10 seconds. **E**. Distribution of firing frequencies on two pGMEAs 60 seconds before (blue) and 60 seconds after (orange) NA stimulation (N = 2 electrodes per device).

We then analyzed the waveforms of the LFP signals recorded from both types of devices. After normalizing the amplitude of the average waveforms recorded through different electrodes, we found that pGMEAs provided signals with similar waveforms with respect to the ones captured using commercial devices. Besides the canonical HL-1 LFP, consisting of a positive peak followed by a negative downward peak and a positive hump, both devices enabled the recording of LFPs with delayed repolarizations and increased initial positive deflections (Figure 4C). These results suggest that, despite having obtained lower amplitude signals, the pGMEAs were able to capture the same types of LFPs as the commercial MEAs.

Finally, we stimulated the cells on two pGMEAs with noradrenaline (NA), in order to determine whether we could capture the activity changes triggered by this compound. After adding NA to a concentration of 0.2 mM, we observed a sustained increase in the frequency of LFP signals (Figure 4D). We quantified the increase of activity over the 90 seconds before and the 90 seconds after stimulation with NA, which moved from an average of 0.44 LFPs/s (σ = 0.08 LFPs/s, N = 2) to 0.81 LFPs/s (σ = 0.09 LFPs/s, N = 2) on one chip and from 0.81 LFPs/s (σ = 0.08 LFPs/s, N = 4) to 1.04 LFPs/s (σ = 0.11 LFPs/s, N = 4), with a significant difference in both cases (Figure 4E).

## Conclusion

This work describes the design and fabrication of printed graphene-based multielectrode arrays and their application for monitoring the electrophysiological activity of cardiac cells. We demonstrated that inkjet printing is a suitable technique for the production of pGMEAs. The dimensions of the features on the printed devices faithfully reproduce the design, and the electrical properties of the electrodes were on par with those of metal- or conductive polymer-based MEAs. The cellular stainings demonstrated that the pGMEAs are a suitable substrate for cardiac cell cultures. The devices could effectively be used to monitor the electrophysiological activity of the cell cultures, confirming both the capabilities of the pGMEAs and their biocompatibility. The main limitations of this proof-of-concept devices are the signal amplitude, which was almost one order of magnitude below that of the signals recorded using commercial MEAs, and the limited throughput of this design. This suggests that modified designs, with a higher number of smaller electrodes, would be a clear way for improveming our pGMEAs. Despite the low amplitudes registered, the LFPs recorded on the printed devices exhibited similar frequencies and waveforms as those on the commercial MEAs. When the cultures were stimulated with noradrenaline, the signals on the pGMEAs effectively reflected the changes in the frequency of LFPs. Therefore, together with the custom signal acquisition system presented here, the pGMEAs we present here can constitute a complete, accessible and fully open-source toolset to perform *in vitro* cardiac electrophysiology.

## Methods

### pGMEA fabrication

#### Graphene Ink preparation

The graphene was produced by electrochemical exfoliation according to the procedure described in a previous publication^66^. Briefly, a piece of graphite foil (Alfa Aesar, 0.13 mm thick, product number: 43078) and a piece of platinum foil, as the anode and cathode respectively, were immersed into 0.1 M (NH_4_)_2_SO^4^ aqueous solution (Millipore, Product Number: 1.01217) at 2 cm. Then a DC voltage of 8 V was applied between two electrodes. Exfoliated graphite flakes were obtained, collected and rinsed through filtration with water and N,N-Dimetylformamid (DMF, Millipore, Product Number : 1.03053) 3 times. Finally, the collection was dispersed in DMF at a concentration of about 2 mg/mL by ultrasonication for 10 minutes.

4 ml electrochemically exfoliated graphene (EEG)/DMF dispersion was centrifuged at 3,000 rpm for 15 min to remove the large flakes. The supernatant was harvested and centrifuged again at 10,500 rpm for 15 minutes to separate EEG from DMF. After the removal of DMF, 40 mg of ethyl cellulose (viscosity 4 cP for 5 w/v% in 80:20 toluene:ethanol, Sigma-Aldrich, product number: 200646) was dispersed in 2 mL 4:1 (volume ratio) mixture solvent of cyclohexanone (Sigma-Aldrich, Product Number: 398241) and terpineol (Sigma-Aldrich, Product Number: 86480), and the dispersion was used to disperse the sedimented EEG by bath ultrasonication for 30 minutes. Finally, the ink was centrifuged at 2,000 rpm for 3 minutes to further remove big particles. The final ink concentration was around 2 mg/mL.

#### Device fabrication

The devices consisting of three layers of different materials were fully printed using a commercial piezoelectric inkjet printer (Dimatix Materials Printer, DMP 2800, Dimatix-Fujifilm Inc.) equipped with 10-pL cartridges (DMC-11610). Layer alignment was done according to the alignment marks in pattern design. First, the commercial silver ink (Advanced Nano Products, Silverjet DGP 40LT-15C) layer was printed with the drop spacing of 40 μm on the glass microscope slides (Marienfeld, 76×52×1 mm, precleaned, reference: 1100420) at room temperature for 2 passes. Then samples were dried on a hot plate at 60°C for 10 minutes and annealed at 120°C for 30 minutes. After cooling down, EEG inks were printed with the drop spacing of 40 μm at the bed temperature of 45°C for 20 passes. Afterward, the samples were dried on a hot plate at 120°C for 10 minutes and annealed in air for 1 h at the temperature around 350°C to remove the ethyl cellulose. Finally, the passivation layer was printed with the same EEG ink but only dried at 120°C to keep the ethyl cellulose. The connector pads around the edge of the devices were covered with silver paste and copper tape to prevent scratches.

### Electrical characterization

The electrochemical system underwent thorough characterization using an AutoLab potentiostat, adhering to the conventional three-electrode setup. For electrochemical impedance spectroscopy, the internal microelectrodes were referenced with an in-planar electrode, complemented by a platinum coil as the counter electrode, all within a 1X PBS electrolyte solution. EIS data were meticulously collected across a frequency range spanning from 1,000,000 Hz to 1 Hz, modulated by a sinusoidal amplitude of 10 mV, while preserving an open circuit potential (OCP).

Images from the full device were taken with a Samsung Galaxy S23 phone camera, and the zoom in of the electrodes by electronic microscopy images were performed using a Zeiss microscope and a 40x, 20x and 10x objectives.

### Cell culture

The HL-1 cardiomyocyte line, derived from the AT-1 mouse atrial cardiomyocyte tumor lineage C57BL/6J), was obtained from Sigma-Aldrich (#ref: SCC065, Missouri, USA). HL-1 cells were cultured in Claycomb medium (#ref: 51800C, Sigma-Aldrich, Missouri, USA) supplemented with 10% fetal bovine serum (#ref: F7524, Sigma-Aldrich, Missouri, USA), 1% L-Glutamine (#ref: 25030-024, Gibco, ThermoFisher Scientific, Massachusetts, USA), 1% penicillin-streptomycin (#ref: P0781, Sigma-Aldrich, Missouri, USA), and 1% norepinephrine (10mM stock) (#ref: A0937-5G, Sigma-Aldrich, Missouri, USA). Cells were seeded onto the pGMEA chips at a density of 25,000 cells/cm^2^ and maintained in an incubator at 37°C in a humidified atmosphere of 5% CO_2_ and 95% air, refreshing the culture media every 24 hours. The chips were previously sterilized by UV irradiation for a period of 3 hours and coated with a solution of 0.0005% fibronectin (#ref: F1141, Sigma-Aldrich, Missouri, USA) and 0.02% gelatin (#ref: G9391-100G, Sigma-Aldrich, Missouri, USA) overnight.

### Cell imaging

To assess biocompatibility, after 48 hours in culture conditions, HL-1 cells were washed three times with 1X PBS and then incubated with MitoTracker Green (#ref M7514, ThermoFisher Scientific, Massachusetts, USA) (diluted in 1X PBS) at a final concentration of 200 nM for 20 minutes at 37°C in the dark. They were then washed three times with 1X PBS and incubated for 10 minutes at 37°C in the dark with Hoechst 33342 (#ref: H21492, ThermoFisher Scientific, Massachusetts, USA) diluted in 1X PBS at a concentration of 1 µg/mL. Afterwards, the cells were washed three times with 1X PBS. The stained cultures were then imaged using the EVOS M5000 Microscope Imaging System (ThermoFisher Scientific, Massachusetts, USA).

### Acquisition system

The signal acquisition system consisted of a PCB board with 2 INTAN Technologies RHD2132 amplifiers connected to a XEM6010 FPGA development which interfaced with a computer via USB connection. A connector board with 64 pogo pins was mounted on the amplifiers’ PCB to interface with the MEAs/pGMEAs. The electrode arrays were held on a custom 3D printed holder. The PCB schematics, lists of components and the MEA holder design are provided as supplementary materials (Supplementary Data 2) and are available online at https://github.com/Leo-GG/pGMEAs/.

### Electrophysiological recordings

The recordings were performed immediately after refreshing the cell culture media. The acquisition was performed using the RHX Data Acquisition Software (INTAN Technologies, USA) with the following parameters: sampling rate = 20 kSamples/s, amplifier lower bandwidth = 0.1 Hz, amplifier upper bandwidth = 100 Hz, notch filter = 50 Hz. We recorded spontaneous activity on two pGMEAs and one commercial MEA for 2 minutes. We then made another set of recordings on HL-1 cultures on 2 pGMEAs, 2 minutes before and 2 minutes after stimulating the cell cultures with 10 µL of 10 mM noradrenaline.

### Data analysis

The signal processing and analysis was performed using the Scipy (version 1.10.1) and Numpy (version 1.24.3) Python libraries. In all cases, we analyzed 90-seconds of data out of the 2 minutes recorded, discarding the beginning and end of each recording. We removed common mode noise by applying a 50 Hz notch filter to the acquired signals. Then we removed low frequency artifacts using a 1Hz high-pass filter. We then computed the 1-second running average of the rectified signal on each channel and labeled all the points with values above 5 times the standard deviation of the rectified, smoothed signal as artifacts, with the aim of removing impulsive noise events. Then we detected putative LFPs by applying the *find_peaks* function from the Scipy library. The amplitude threshold for peak detection was 3 times the standard deviation of the recording on each channel, excluding all values labeled as artifacts, and the minimum distance between peaks was set to 0.25 seconds (5000 samples). We considered the 0.5 seconds around each detected peak as the corresponding waveform of the putative LFP. To ensure that the signals corresponded to actual LFPs, we kept only the ones with homogeneous waveforms. To do so, we computed the RMSD of each waveform, and we removed those with a z-score above 2.

