## Supplementary figures and images for "Inkjet-printed graphene multielectrode arrays: an accessible platform for *in vitro* cardiac electrophysiology"

### Ag 40um.bmp

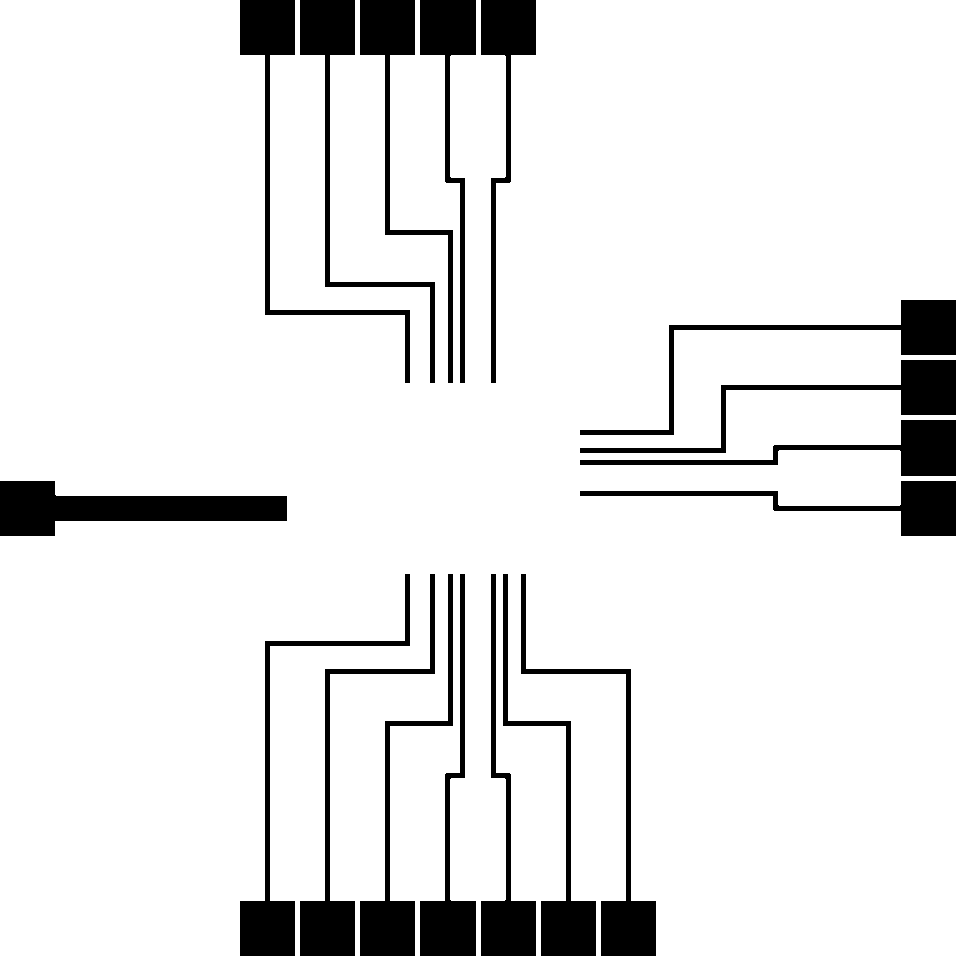

### central graphene 40um.bmp

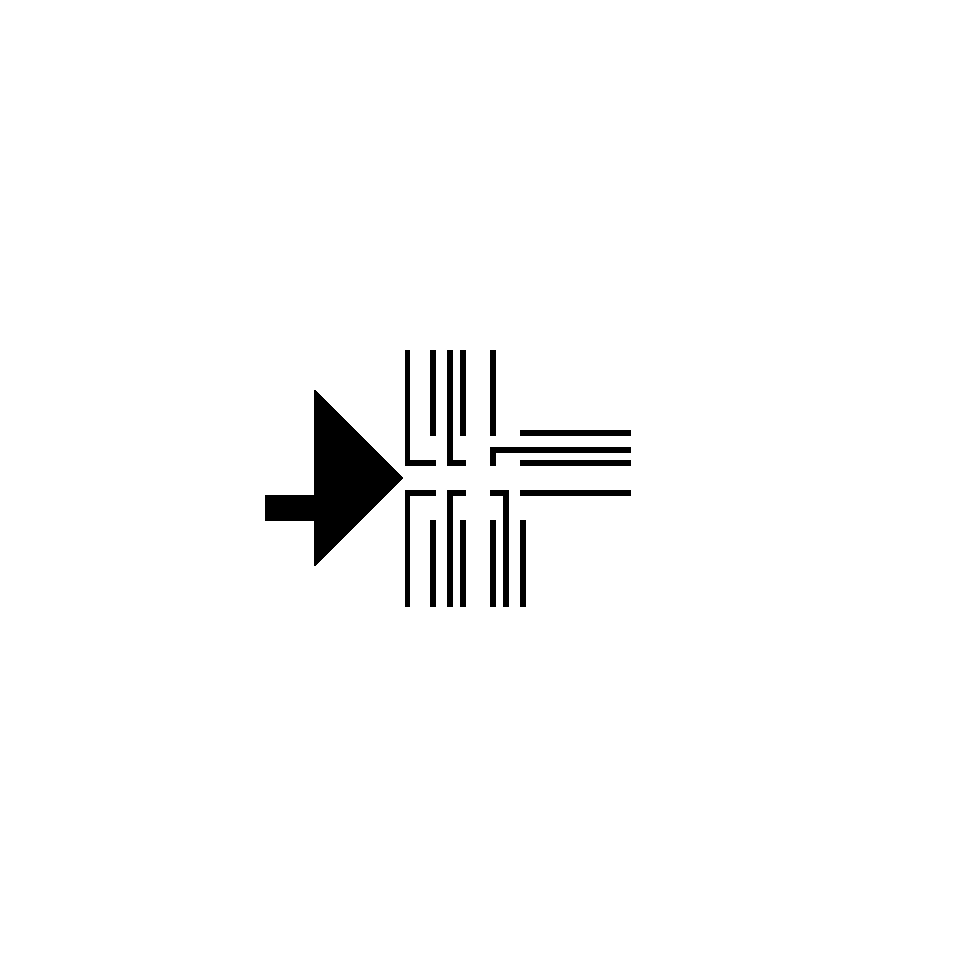

### passivation 40um.bmp

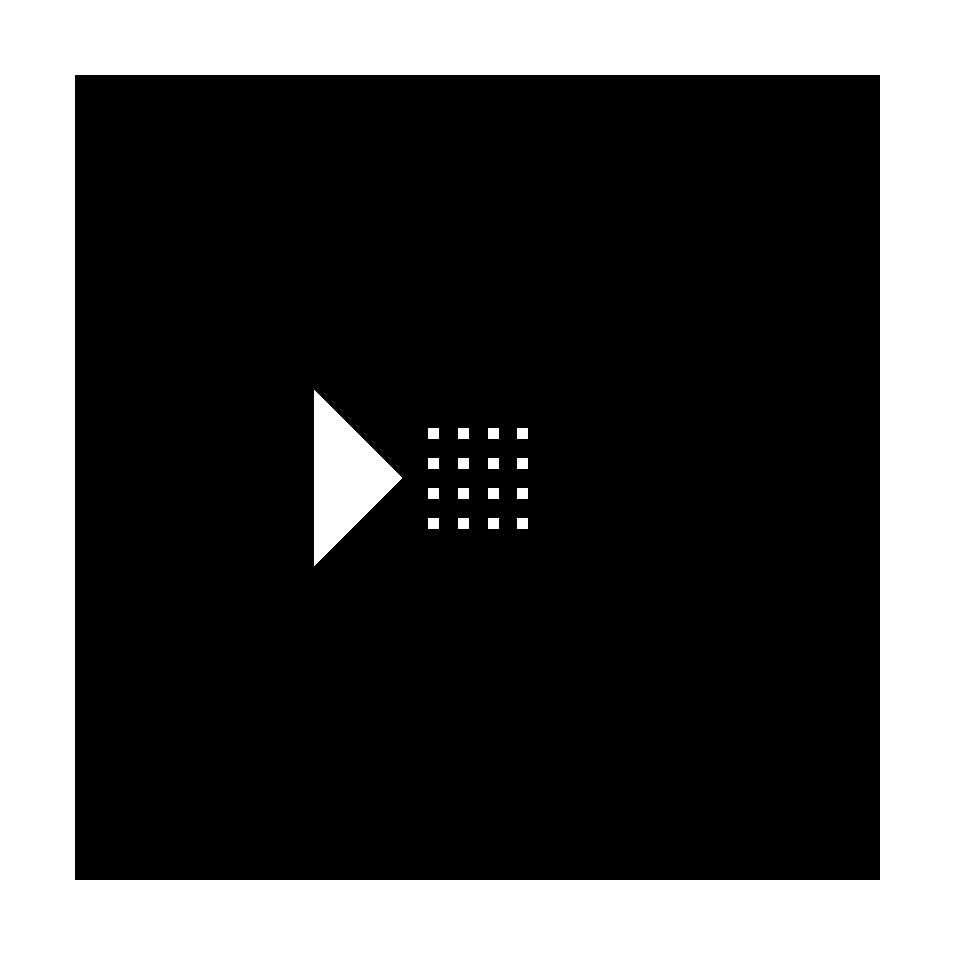
