## Supplementary data 2 for "Inkjet-printed graphene multielectrode arrays: an accessible platform for *in vitro* cardiac electrophysiology": amp_boardV3.pro

```
update=10/16/21 13:41:19
version=1
last_client=pcbnew
[general]
version=1
RootSch=
BoardNm=
[cvpcb]
version=1
NetIExt=net
[eeschema]
version=1
LibDir=
[eeschema/libraries]
[schematic_editor]
version=1
PageLayoutDescrFile=
PlotDirectoryName=
SubpartIdSeparator=0
SubpartFirstId=65
NetFmtName=Pcbnew
SpiceAjustPassiveValues=0
LabSize=50
ERC_TestSimilarLabels=1
[pcbnew]
version=1
PageLayoutDescrFile=
LastNetListRead=amp_boardV3.net
CopperLayerCount=4
BoardThickness=1.6
AllowMicroVias=0
AllowBlindVias=0
RequireCourtyardDefinitions=0
ProhibitOverlappingCourtyards=1
MinTrackWidth=0.2
MinViaDiameter=0.4
MinViaDrill=0.3
MinMicroViaDiameter=0.2
MinMicroViaDrill=0.09999999999999999
MinHoleToHole=0.25
TrackWidth1=0.25
TrackWidth2=0.5
ViaDiameter1=0.8
ViaDrill1=0.4
dPairWidth1=0.2
dPairGap1=0.25
dPairViaGap1=0.25
SilkLineWidth=0.15
SilkTextSizeV=1
SilkTextSizeH=1
SilkTextSizeThickness=0.15
SilkTextItalic=0
SilkTextUpright=1
CopperLineWidth=0.2
CopperTextSizeV=1.5
CopperTextSizeH=1.5
CopperTextThickness=0.3
CopperTextItalic=0
CopperTextUpright=1
EdgeCutLineWidth=0.15
CourtyardLineWidth=0.05
OthersLineWidth=0.15
OthersTextSizeV=1
OthersTextSizeH=1
OthersTextSizeThickness=0.15
OthersTextItalic=0
OthersTextUpright=1
SolderMaskClearance=0.051
SolderMaskMinWidth=0.25
SolderPasteClearance=0
SolderPasteRatio=-0
[pcbnew/Layer.F.Cu]
Name=F.Cu
Type=0
Enabled=1
[pcbnew/Layer.In1.Cu]
Name=In1.Cu
Type=0
Enabled=1
[pcbnew/Layer.In2.Cu]
Name=In2.Cu
Type=0
Enabled=1
[pcbnew/Layer.In3.Cu]
Name=In3.Cu
Type=0
Enabled=0
[pcbnew/Layer.In4.Cu]
Name=In4.Cu
Type=0
Enabled=0
[pcbnew/Layer.In5.Cu]
Name=In5.Cu
Type=0
Enabled=0
[pcbnew/Layer.In6.Cu]
Name=In6.Cu
Type=0
Enabled=0
[pcbnew/Layer.In7.Cu]
Name=In7.Cu
Type=0
Enabled=0
[pcbnew/Layer.In8.Cu]
Name=In8.Cu
Type=0
Enabled=0
[pcbnew/Layer.In9.Cu]
Name=In9.Cu
Type=0
Enabled=0
[pcbnew/Layer.In10.Cu]
Name=In10.Cu
Type=0
Enabled=0
[pcbnew/Layer.In11.Cu]
Name=In11.Cu
Type=0
Enabled=0
[pcbnew/Layer.In12.Cu]
Name=In12.Cu
Type=0
Enabled=0
[pcbnew/Layer.In13.Cu]
Name=In13.Cu
Type=0
Enabled=0
[pcbnew/Layer.In14.Cu]
Name=In14.Cu
Type=0
Enabled=0
[pcbnew/Layer.In15.Cu]
Name=In15.Cu
Type=0
Enabled=0
[pcbnew/Layer.In16.Cu]
Name=In16.Cu
Type=0
Enabled=0
[pcbnew/Layer.In17.Cu]
Name=In17.Cu
Type=0
Enabled=0
[pcbnew/Layer.In18.Cu]
Name=In18.Cu
Type=0
Enabled=0
[pcbnew/Layer.In19.Cu]
Name=In19.Cu
Type=0
Enabled=0
[pcbnew/Layer.In20.Cu]
Name=In20.Cu
Type=0
Enabled=0
[pcbnew/Layer.In21.Cu]
Name=In21.Cu
Type=0
Enabled=0
[pcbnew/Layer.In22.Cu]
Name=In22.Cu
Type=0
Enabled=0
[pcbnew/Layer.In23.Cu]
Name=In23.Cu
Type=0
Enabled=0
[pcbnew/Layer.In24.Cu]
Name=In24.Cu
Type=0
Enabled=0
[pcbnew/Layer.In25.Cu]
Name=In25.Cu
Type=0
Enabled=0
[pcbnew/Layer.In26.Cu]
Name=In26.Cu
Type=0
Enabled=0
[pcbnew/Layer.In27.Cu]
Name=In27.Cu
Type=0
Enabled=0
[pcbnew/Layer.In28.Cu]
Name=In28.Cu
Type=0
Enabled=0
[pcbnew/Layer.In29.Cu]
Name=In29.Cu
Type=0
Enabled=0
[pcbnew/Layer.In30.Cu]
Name=In30.Cu
Type=0
Enabled=0
[pcbnew/Layer.B.Cu]
Name=B.Cu
Type=0
Enabled=1
[pcbnew/Layer.B.Adhes]
Enabled=1
[pcbnew/Layer.F.Adhes]
Enabled=1
[pcbnew/Layer.B.Paste]
Enabled=1
[pcbnew/Layer.F.Paste]
Enabled=1
[pcbnew/Layer.B.SilkS]
Enabled=1
[pcbnew/Layer.F.SilkS]
Enabled=1
[pcbnew/Layer.B.Mask]
Enabled=1
[pcbnew/Layer.F.Mask]
Enabled=1
[pcbnew/Layer.Dwgs.User]
Enabled=1
[pcbnew/Layer.Cmts.User]
Enabled=1
[pcbnew/Layer.Eco1.User]
Enabled=1
[pcbnew/Layer.Eco2.User]
Enabled=1
[pcbnew/Layer.Edge.Cuts]
Enabled=1
[pcbnew/Layer.Margin]
Enabled=1
[pcbnew/Layer.B.CrtYd]
Enabled=1
[pcbnew/Layer.F.CrtYd]
Enabled=1
[pcbnew/Layer.B.Fab]
Enabled=1
[pcbnew/Layer.F.Fab]
Enabled=1
[pcbnew/Layer.Rescue]
Enabled=0
[pcbnew/Netclasses]
[pcbnew/Netclasses/Default]
Name=Default
Clearance=0.2
TrackWidth=0.25
ViaDiameter=0.8
ViaDrill=0.4
uViaDiameter=0.3
uViaDrill=0.1
dPairWidth=0.2
dPairGap=0.25
dPairViaGap=0.25
[pcbnew/Netclasses/1]
Name=Pour
Clearance=0.2
TrackWidth=0.5
ViaDiameter=0.8
ViaDrill=0.4
uViaDiameter=0.3
uViaDrill=0.1
dPairWidth=0.2
dPairGap=0.25
dPairViaGap=0.25
```
