## Supplementary figures 1 and 2 for "Inkjet-printed graphene multielectrode arrays: an accessible platform for *in vitro* cardiac electrophysiology"

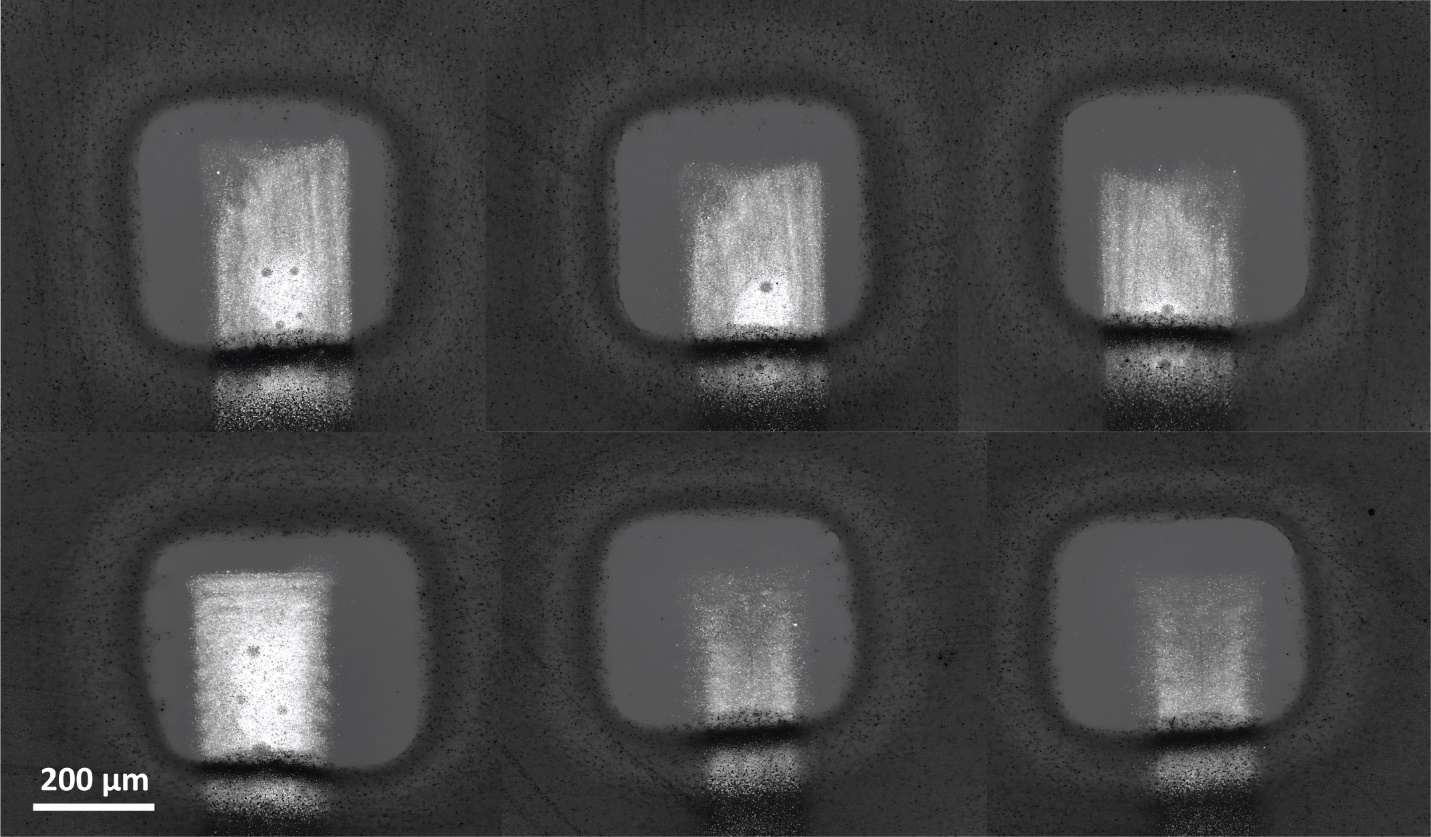

**Figure S1**. Light-saturated micrographs from electrode openings showing details from inkjet printed electrodes.

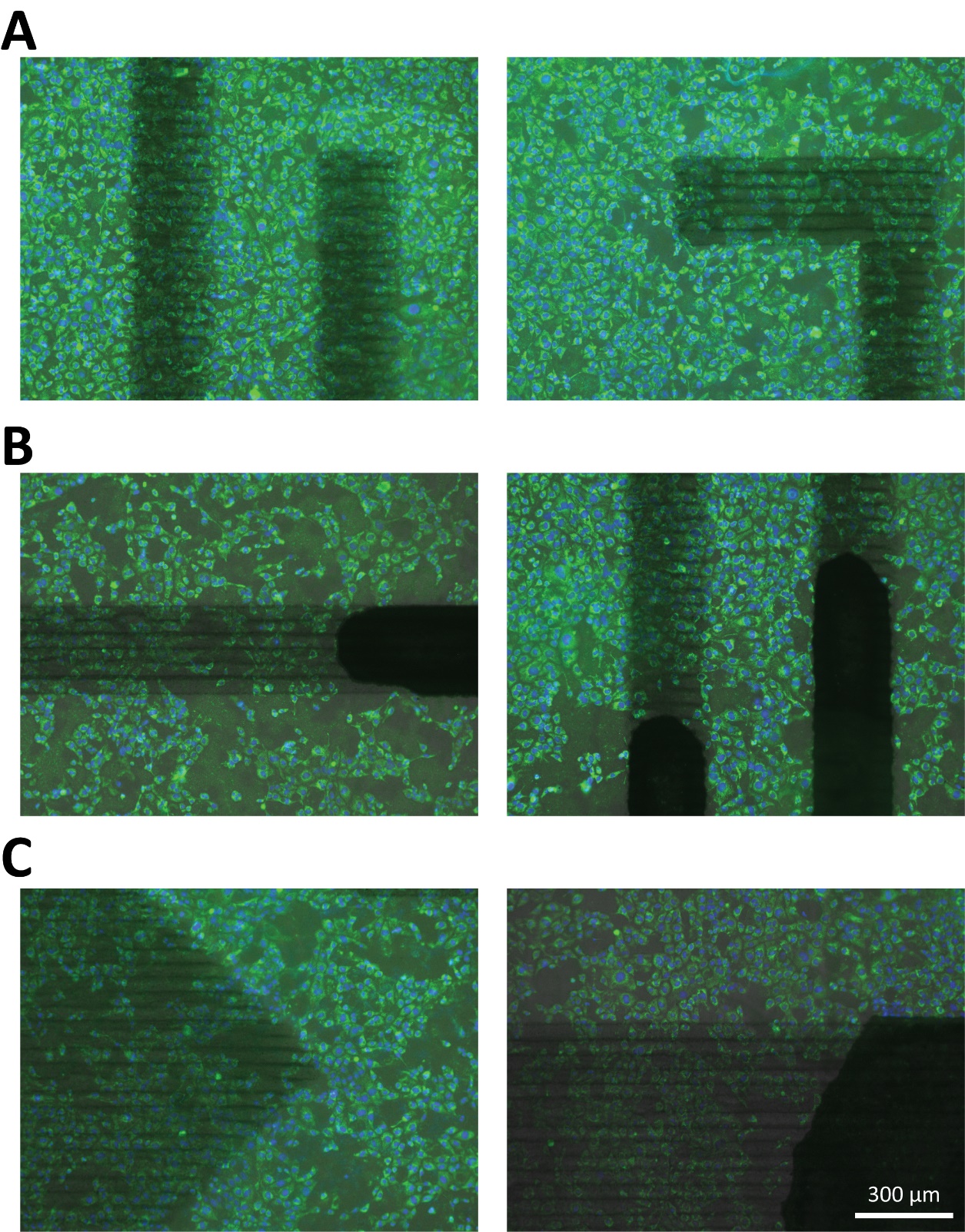

**Figure S2**. MitoTracker Green (green) and Hoechst 33342 (blue) stainings of HL-1 cells plated on pGMEA. Cell can be seen growing around all the elements present in the devices: the exposed annealed graphene electrodes (**A**), the graphene and silver traces (**B**) and the reference electrode (**C**).
